# Genetic and Behavioral Variation Underlying an Integrated Temporal Phenotype in an Admixed South American Population

**DOI:** 10.64898/2026.08.18.745563

**Authors:** Mariana Marchesano, Lucía Spangenberg, Cecilia Casaravilla, Julieta Castillo, Ana Silva, Bettina Tassino

## Abstract

Chronotype is a complex trait reflecting individual differences in the temporal organization of rest and activity, with important health implications. The Uruguayan population, characterized by a tri-hybrid origin (African, European, and Indigenous), exhibits a bias toward eveningness. The genetic variation in clock genes underlying chronotype in this population remains unexplored. To address this gap, we analyze healthy young adults from the extremes of the chronotype distribution (early, *n* = 37; late, *n* = 38; 63% female; 23.1 ± 3.4 years), integrating self-reported measures, actigraphy, and low-pass whole-genome sequencing. Global ancestry is predominantly European, with Indigenous and African components, and does not differ between chronotypes. Variant density is highest in *PER2*. T-allele carriers of a *PER2* variant previously associated with late chronotypes (rs35333999) differ from non-carriers in activity acrophase. Multidimensional scaling of variants across 19 canonical clock genes reveal differential representation of early and late chronotypes across genetic clusters. When examined by functional groups, the signal is restricted to genes involved in degradation of the circadian clock’s repressor arm, with *BTRC*, a mediator of PER2 degradation, showing the same pattern when assessed individually. We derive a joint behavioral component capturing the variation in food intake, moderate-to-vigorous physical activity, light exposure, and sleep timing, which correlates with dim-light melatonin onset (DLMO), the gold-standard marker of circadian phase, and show differences among genetic clusters. Our integrative multilevel approach suggests a complex interplay between behavioral and genetic factors shaping chronotype in this cohort, highlighting the *PER2–BTRC* axis as a candidate mechanism for future investigations.

## INTRODUCTION

Virtually all living organisms follow internal rhythms that are in turn guided by environmental cues (Paranjpe and Sharma 2005). Circadian rhythms are self-sustained oscillations with a period of ∼24 hours that respond primarily to the light/dark cycle, but are also influenced by other time givers, or zeitgebers, such as food intake and physical activity (Albrecht 2010; Mistlberger and Skene 2005; Youngstedt et al. 2019). At a physiological level, the circadian system consists of a network of oscillators. The suprachiasmatic nucleus of the hypothalamus (SCN) receives photic input from intrinsically photosensitive retinal ganglion cells and establishes an internal rhythmic environment that entrains peripheral oscillators throughout the body (Roenneberg et al. 2022). The pineal hormone melatonin, a biological time-domain–acting molecule, is synthesized under the control of the SCN and tightly coupled to the light/dark cycle, with production occurring during the night and being suppressed by light exposure (Arendt 2005; Cipolla-Neto and Amaral 2018). The onset of melatonin secretion under dim-light conditions (DLMO) is the gold-standard biomarker of circadian phase (Pandi-Perumal et al. 2007).

An individual’s circadian timing of rest and activity is commonly referred to as chronotype (Roenneberg et al. 2003). Earlier chronotypes (colloquially, “larks”) exhibit earlier circadian phases than later chronotypes (“owls”) and have been associated with shorter circadian periods (Duffy et al. 2001; Brown et al. 2008). Although grounded in circadian systems biology, this temporal phenotype is also shaped by cultural and geographic contexts (Leocadio-Miguel et al. 2017; Golombek 2026). Compared with rural populations, urban residents exhibit later chronotypes and lower daytime light exposure (Carvalho et al. 2014). Artificial light at night delays circadian phase (Wright et al. 2013; Moreno et al. 2015; Casiraghi et al. 2020).

Circadian rhythms are underpinned by highly conserved genetic mechanisms (Von Schantz et al. 2021). Clock genes were among the first genes linked to a behavioral phenotype (Konopka and Benzer 1971), and circadian clock proteins have since been implicated in diverse physiological and health-related processes, including metabolism, immune function, cancer, and neurodegeneration (Takahashi 2021). The core clock mechanism relies on a transcriptional-translational feedback loop (TTFL) in which translated proteins regulate their own transcription (Cox and Takahashi 2019; Patton and Hastings 2023). Circadian phenotyping based on the expression of core-clock genes has recently been proposed as an alternative approach (Dose et al. 2023). However, the implementation of molecular phenotyping strategies requires validation across diverse populations, as these models are population-dependent due to differences in genomic background and allele frequencies (Biscontin et al. 2025).

Genome-wide association studies (GWAS) have identified single nucleotide polymorphism (SNPs) associated with morningness; however, most studies have been conducted in homogeneous adult European populations (Jones et al. 2019a; Lane et al. 2016, 2023; Paz et al. 2023). PER2 variant rs35333999 has been previously associated with longer period and late chronotypes (Ashbrook et al. 2020; Vera et al. 2018), and reported to be present only in populations with known European and Indigenous admixture (Emmanuel and von Schantz 2018). Chang et al. (2019) analysed this variant in a large UK Biobank cohort and in a multiethnic sample, reporting a significant association with longer intrinsic circadian periods and suggesting a biological mechanism for inter-individual differences in chronotype.

Studies in an admixed Brazilian population have reported an association between Indigenous (but not African or European) ancestry and morning preference (Egan et al. 2017; Von Schantz et al. 2015). The Uruguayan population presents an admixed genomic landscape composed mainly of European components (∼78%), with smaller proportions of Indigenous (∼14%) and African components (∼8%), as a result of European invasions and successive waves of migration, indigenous genocide, and the enslavement of African people (Sans et al. 1997; Spangenberg et al. 2021 and unpublished data). Interestingly, the Uruguayan population has shown a tendency towards eveningness, as observed across different cohorts and ages with varying exposure to light, exercise schedules, and social pressures (Tassino and Leone 2025). However, genetic data on circadian traits have not yet been reported in this population.

This is the first study to comprehensively characterize chronotype in this underrepresented admixed population by integrating behavioral, physiological, and genomic data.

## RESULTS

We report data from a young Uruguayan cohort assembled to investigate the genetic and behavioral factors associated with chronotype, providing the first characterization of variation in clock genes in this population. To maximize phenotypic contrast, participants were selected from the upper and lower quartiles of the chronotype distribution, defined by the mid-sleep point on free days corrected for sleep debt on weekdays (MSFsc; Roenneberg et al. 2004). Fig. 1 summarizes cohort assembly, including participant selection and quality-control procedures, and illustrates the distribution of chronotype highlighting the individuals selected for the analysis. Early (n = 37; MSFsc = 03:18 ± 00:39 h) and late chronotypes (n = 38; MSFsc = 06:59 ± 00:48 h) did not differ in age, sex or BMI (23.4 ± 3.2 vs. 22.9 ± 3.6 years, 59% vs. 66% female, 24.5 ± 2.9 vs. 24.7 ± 3.9 kg/m²; all P > 0.9).

**Figure 1.**
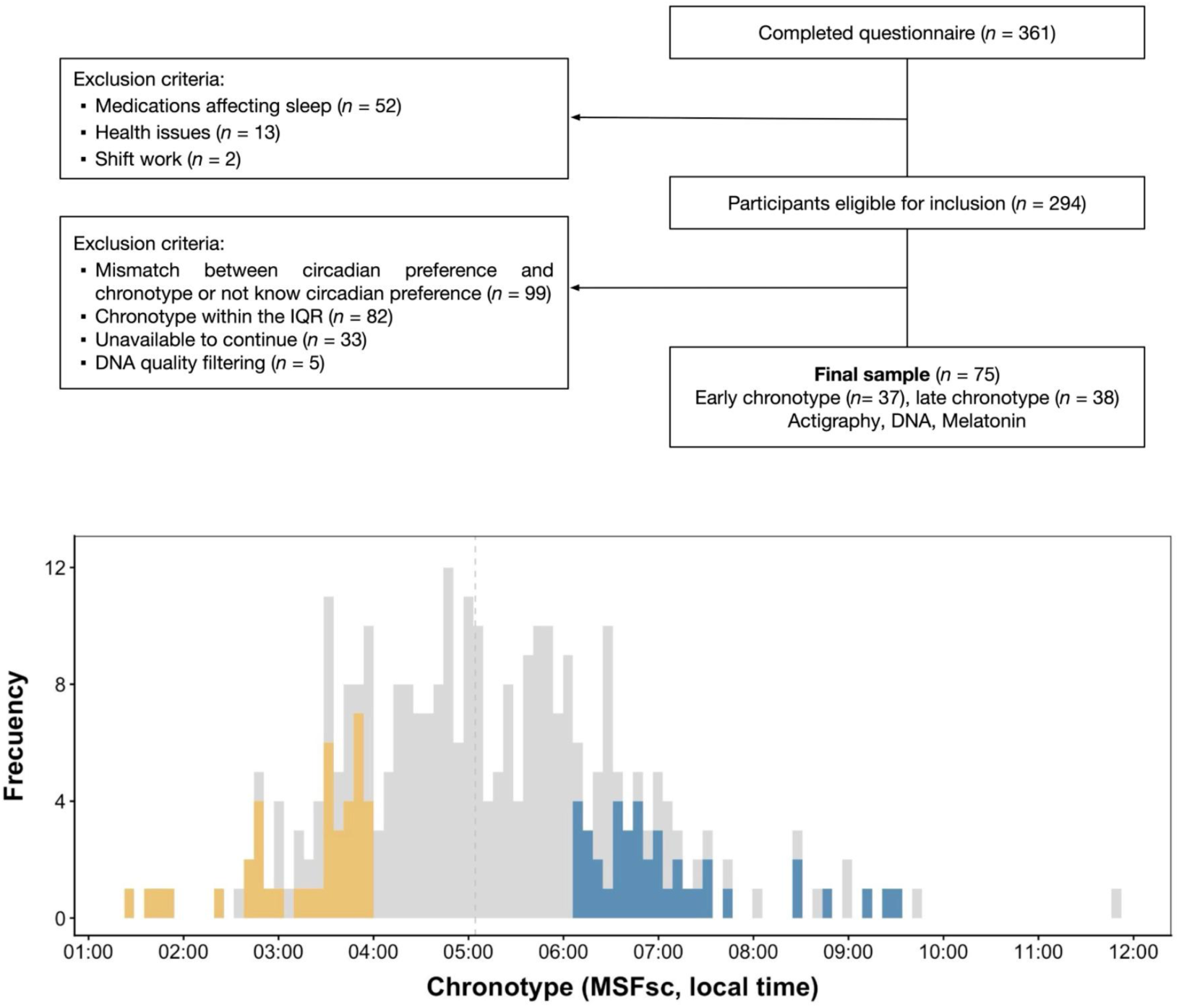
Top, flow chart of study participation. Bottom, chronotype histogram showing the distribution of the full sample, with participants who continued in the study highlighted in color (yellow, early chronotype; blue, late chronotype).

### Admixture structure in the cohort

Given the admixed background of the Uruguayan population, we first characterized the global ancestry composition of the cohort using whole-genome sequencing. Ancestry proportions were 82.98 ± 12.3% European (range 47.8–100%), 7.46 ± 8.1 % Indigenous (range 0.0–33.4%), 7.36 ± 6.6% African (range 0.0–38.1%), and 2.2% Other (Asia and Oceania). Southwest Europe represented the largest contribution (38.35%), followed by Eastern Mediterranean (13.89%), and Northern Italy (11.99%). Individual ancestry profiles based on subcontinental estimates are illustrated in Fig. 2A. Global ancestry proportions did not differ between early and late chronotypes (P > 0.9). Likewise, chronotype was not associated with the first four genome-wide multidimensional scaling (MDS) components after adjusting for sex in a logistic regression model (all P > 0.1). Genome-wide population structure and its relationship with global ancestry proportions and chronotype are shown in Fig. S1.

**Figure 2.**
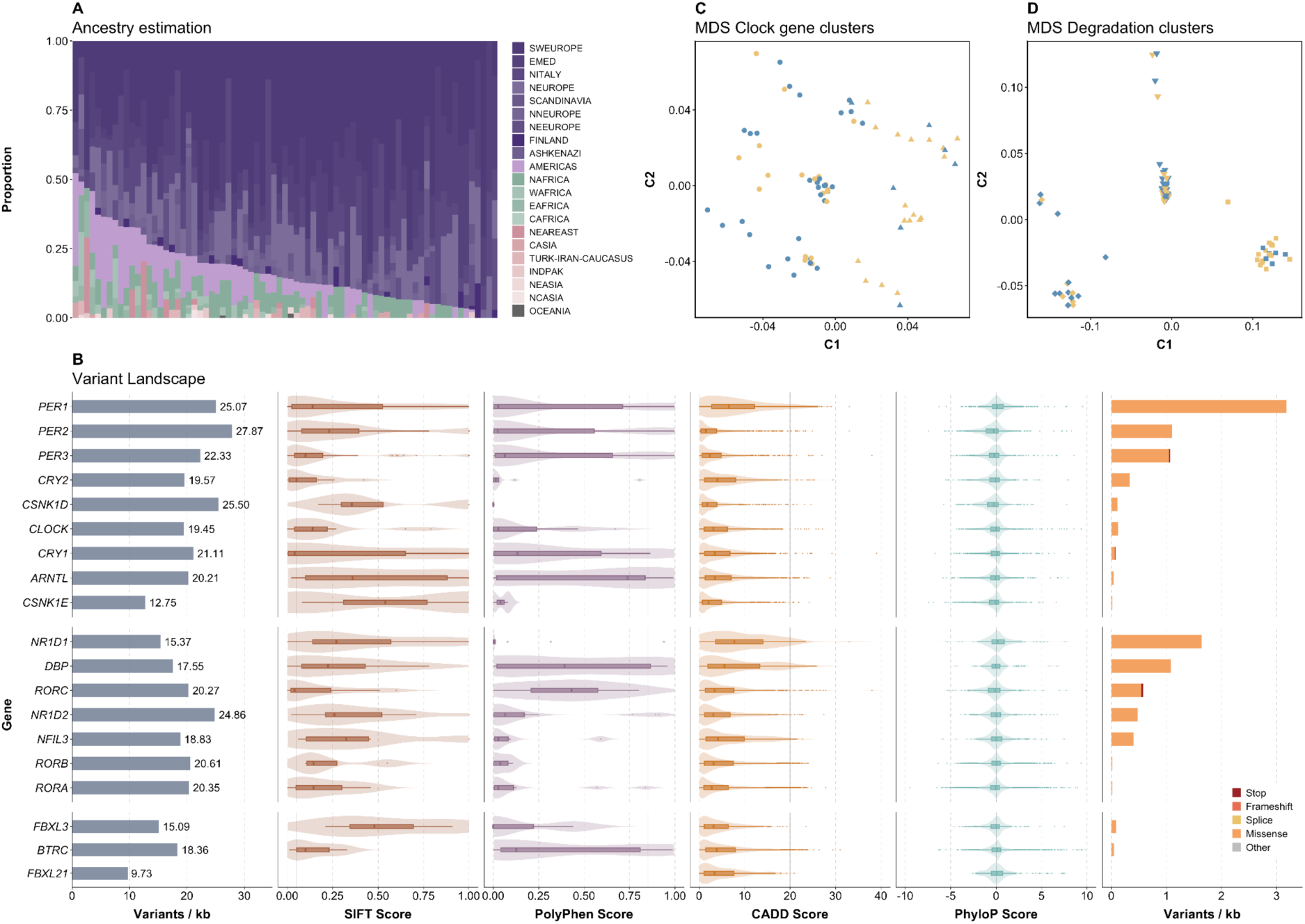
A, admixture plot showing ancestry proportions relative to the GENCOVE reference panel. B, variant density (variants per kb, normalized by gene length), in silico pathogenicity scores (SIFT, PolyPhen, and CADD), conservation scores (PhyloP), and variant consequences (stop-gained, frameshift, splice-site, and missense), normalized by gene length. Solid gray lines indicate score thresholds. C, multidimensional scaling (MDS) plots for clock genes. D, MDS plots for genes involved in the degradation pathway, including *FBXL3*, *FBXL21*, and *BTRC*. Clusters are represented by shape, and chronotypes by color (yellow, early; blue, late).

### Clock gene variant landscape

We analyzed 38239 SNPs across 19 genes previously implicated in the circadian clock (Fig. S2), including 296 missense variants and 4 stop-gained variants (Fig. 2B). Variant density varied across genes, with *PER2*, *CSNK1D*, and *PER1* showing the highest number of variants per kilobase (27.87, 25.5, and 25.07 respectively) (Table S1). The maximum predicted deleteriousness CADD scores were observed for variants in *CRY1* (39), *PER3* and *RORC* (38), *NR1D1*, *PER1* and *PER2* (33), and *BTRC* (31) (Table S1). Evolutionary conservation (PhyloP) was higher for variants in *NR1D1* (mean = 0.35) and lower for *PER2* (mean = −0.47), with maximum values in *RORC, BTRC, PER1, CLOCK y RORB* (> 9), and minimum in *RORA*, *CSNK1E* and *PER2* (< −7). Among missense variants, 23% were predicted deleterious by SIFT, and 10% were predicted probably damaging by PolyPhen (Table S2).

The number of observed variants per individual (with at least one alternative allele) was 739 ± 118 (range, 531–932) in early chronotypes and 801 ± 120 (range, 498–1023) in late chronotypes. The proportion of variants with CADD > 20 (a common threshold for potentially pathogenic variants, since it corresponds to the top 1% of most deleterious variants) among observed variants per individual showed significant differences between chronotypes (P = 0.006), with slightly higher median proportion in early chronotypes (early: 0.003; late: 0.002, r = 0.32). At the gene level, CADD >20 variants were observed across a subset of clock genes, with *RORA* and *BTRC* accounting for the largest numbers (41 and 23 variants in early chronotypes and 31 and 21 in late chronotypes, respectively). No significant differences were observed for the other in silico annotations, including SIFT, PolyPhen, and PhyloP.

Genotyping of the *PER2* variant rs35333999, previously associated with later chronotype, identified eight T-allele carriers (TC, six late and two early chronotypes) and 67 homozygous non-carriers (CC, 32 late and 35 early chronotypes). No significant differences were found between T-allele carriers and non-carriers for categorical chronotype.

### Clock gene clusters and chronotype

Following LD-pruning and additional quality control, 2338 independent variants were retained for multidimensional scaling (MDS). Hierarchical clustering on MDS components identified two clusters when the full clock gene set was considered, although cluster separation was weak (silhouette score = 0.27) (Fig. 2C). Three clusters were identified for the core gene set and two for the auxiliary loop gene set, both with weak separation (silhouette scores 0.36 and 0.22, respectively). In contrast, the proteasomal degradation pathway gene set showed clear cluster separation (silhouette score = 0.65) (Fig. 2D).

Comparison of cluster assignments across chronotypes revealed significant differences for the full clock gene set, with a large effect size (Cramér’s V in Table 1). At the functional level, no significant differences were observed for the core clock or auxiliary loop gene sets. The proteasomal degradation pathway gene set showed a significant association with chronotype. However, this association did not remain significant after correction for multiple testing, and was characterized by a modest effect size (Table 1).

**Table 1.** Chronotype comparison among clock gene clusters.

|  | Chronotype |  | P | P adj. | Cramér's V |
| --- | --- | --- | --- | --- | --- |
|  | Early (*n* = 37) | Late (*n* = 38) |  |  |  |
| Clock genes full set (19 genes) |  |  | **0.007** | **0.029** | 0.3 |
| Cluster 1 | 51% (19) | 82% (31) |  |  |  |
| Cluster 2 | 49% (18) | 18% (7) |  |  |  |
| Core TTFL set (9 genes) |  |  | 0.6 | >0.9 | 0 |
| Cluster 1 | 49% (18) | 39% (15) |  |  |  |
| Cluster 2 | 30% (11) | 29% (11) |  |  |  |
| Cluster 3 | 22% (8) | 32% (12) |  |  |  |
| Auxiliary loop set (7 genes) |  |  | >0.9 | >0.9 | 0 |
| Cluster 1 | 35% (13) | 37% (14) |  |  |  |
| Cluster 2 | 65% (24) | 63% (24) |  |  |  |
| Degradation pathway set (3 genes) |  |  | **0.03** | 0.09 | 0.26 |
| Cluster 1 | 41% (15) | 55% (21) |  |  |  |
| Cluster 2 | 16% (6) | 29% (11) |  |  |  |
| Cluster 3 | 43% (16) | 16% (6) |  |  |  |

% (n); Fisher's exact test. Holm correction for multiple testing. Clock gene clusters were detected by MDS on variants from *BMAL1*, *CLOCK*, *PER*, *CRY*, *CK1* (Core TTFL), *ROR*, *REV-ERB*, *NFIL3*, *DBP* (Auxiliary loop), *FBXL3*, *FBXL21*, *BTRC* (Degradation pathway).

### Gene-level decomposition of the degradation pathway signal

Of the three genes comprising the degradation pathway set (*BTRC*, *FBXL3*, and *FBXL21*), only *BTRC* reproduced a significant association with chronotype, showing a medium effect size (Fisher’s exact P = 0.03, Cramér’s V = 0.26) and a cluster silhouette (0.73) even higher than that of the combined gene set (0.65). *FBXL3* and *FBXL21* showed no association with chronotype (both P > 0.05, cluster silhouette 0.7 and 0.40 respectively).

### Circadian phase and behavioral timing

The gold-standard marker of circadian phase, DLMO, was delayed by 1 h 54 min in late compared with early chronotypes, providing physiological validation of the chronotype classification based on self-reported measures (Table 2). Actigraphy-derived circadian phase indicators showed a similar pattern with late chronotypes consistently showing later rest–activity timing by 1 h 40 min in midpoint of the most active 10 h (M10c), by 1 h 42 min in the midpoint of the least active 5 h (L5c). The largest difference was observed for acrophase estimated by cosinor model, which was delayed by 2 h 02 min in late compared with early chronotypes.

**Table 2.**
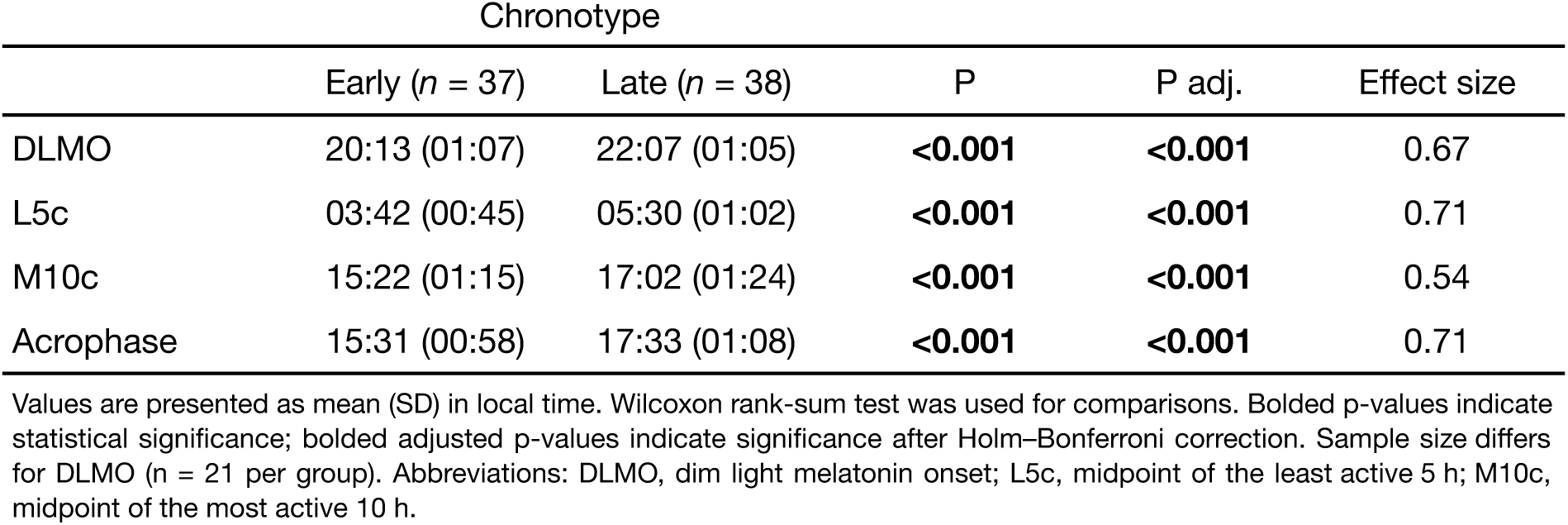
Circadian phase indicators.

|  | Chronotype |  | P | P adj. | Effect size |
| --- | --- | --- | --- | --- | --- |
| | Early ($n = 37$) | Late ($n = 38$) | | | |
| DLMO | 20:13 (01:07) | 22:07 (01:05) | **<0.001** | **<0.001** | 0.67 |
| L5c | 03:42 (00:45) | 05:30 (01:02) | **<0.001** | **<0.001** | 0.71 |
| M10c | 15:22 (01:15) | 17:02 (01:24) | **<0.001** | **<0.001** | 0.54 |
| Acrophase | 15:31 (00:58) | 17:33 (01:08) | **<0.001** | **<0.001** | 0.71 |

Values are presented as mean (SD) in local time. Wilcoxon rank-sum test was used for comparisons. Bolded p-values indicate statistical significance; bolded adjusted p-values indicate significance after Holm–Bonferroni correction. Sample size differs for DLMO ( $n = 21$ per group). Abbreviations: DLMO, dim light melatonin onset; L5c, midpoint of the least active 5 h; M10c, midpoint of the most active 10 h.

To further examine the genetic contribution to circadian phase, we focused on the *PER2* rs35333999 variant, which showed a consistent directional pattern across all four phase indicators. T-allele carriers exhibited later timing for DLMO (1 h 04 min), L5c (38 min), M10c (55 min), and acrophase (1 h 08 min) compared with non-carriers (Table S3). The largest difference was again observed for acrophase, which showed a significant difference between genotype groups (P = 0.04, r = 0.24), although this association did not withstand Holm–Bonferroni correction.

To characterize the temporal organization of daily behaviors, we assessed the timing of food intake, physical activity, light exposure, and sleep. Specifically, we evaluated first and last food intake times obtained by self-report and actigraphy-derived first and last times of physical activity above the moderate-to-vigorous physical activity (MVPA) threshold, first and last times of light exposure above an illuminance threshold of 100 lux, sleep onset and offset times. Late chronotypes exhibited significant delays across all behavioral timing indicators after adjustment for multiple comparisons (all adjusted P < 0.001), with effect sizes ranging from medium (last food consumption on weekdays) to large (all remaining) (Table 3). Among habitual coffee and yerba mate infusion consumers, late chronotypes showed later timing of daily consumption, particularly for first coffee and yerba mate intake on weekends, with moderate-to-large effect sizes (Table S4).

**Table 3.** Behavioral timing by chronotype.

|  | Chronotype |  | P | P adj. | Effect size |
| --- | --- | --- | --- | --- | --- |
| | Early ($n = 37$) | Late ($n = 38$) | | | |
| **Food consumption** |  |  |  |  |  |
| First weekdays | 08:22 (02:34) | 10:44 (01:53) | **<0.001** | **<0.001** | 0.58 |
| Last weekdays | 21:39 (01:08) | 22:39 (01:05) | **<0.001** | **<0.001** | 0.41 |
| First weekends | 09:44 (01:30) | 12:36 (01:26) | **<0.001** | **<0.001** | 0.73 |
| Last weekends | 21:47 (01:04) | 23:34 (01:20) | **<0.001** | **<0.001** | 0.61 |
| **Moderate-to-vigorous physical activity** |  |  |  |  |  |
| First weekdays | 07:44 (00:58) | 10:02 (01:26) | **<0.001** | **<0.001** | 0.69 |
| Last weekdays | 23:11 (01:04) | 01:06 (01:07) | **<0.001** | **<0.001** | 0.69 |
| First weekends | 08:57 (01:02) | 11:07 (01:18) | **<0.001** | **<0.001** | 0.69 |
| Last weekends | 23:52 (01:23) | 01:37 (01:27) | **<0.001** | **<0.001** | 0.53 |
| **Light exposure** |  |  |  |  |  |
| First weekdays | 07:18 (00:50) | 09:28 (01:14) | **<0.001** | **<0.001** | 0.73 |
| Last weekdays | 23:14 (01:13) | 01:08 (01:08) | **<0.001** | **<0.001** | 0.66 |
| First weekends | 08:24 (00:57) | 10:35 (01:15) | **<0.001** | **<0.001** | 0.71 |
| Last weekends | 00:03 (01:30) | 01:41 (01:28) | **<0.001** | **<0.001** | 0.53 |
| **Sleep** |  |  |  |  |  |
| Sleep end weekdays | 07:10 (00:56) | 09:20 (01:15) | **<0.001** | **<0.001** | 0.7 |
| Sleep onset weekdays | 23:45 (01:04) | 01:46 (01:05) | **<0.001** | **<0.001** | 0.71 |
| Sleep end weekends | 08:17 (00:58) | 10:29 (01:17) | **<0.001** | **<0.001** | 0.71 |
| Sleep onset weekends | 00:46 (01:20) | 02:36 (01:29) | **<0.001** | **<0.001** | 0.59 |

Values are given as mean (SD) in local time. Wilcoxon rank-sum test. Bold-formatted p-values indicate statistical significance. Bold-formatted adjusted p-value indicate statistical significance after Holm–Bonferroni correction.

### An integrated temporal phenotype

To identify a shared variation across temporal organization across behavioral domains and account for their correlated structure, we applied Joint and Individual Variation Explained (JIVE), a multivariate method that decomposes variation into joint, individual, and residual components. We analyzed four domains—food intake, MVPA, light exposure, and sleep—each represented by four timing variables corresponding to the first and last events on weekdays and weekends. JIVE identified a single joint component capturing the shared temporal structure across all four domains, together with domain-specific individual components (food: 1, MVPA: 1, light: 2, sleep: 1). The joint component explained for 39.6%, 75.2%, 74.5%, and 75.4% of the variance in the food, MVPA, light, and sleep domains, respectively (Fig. 3A), indicating that a substantial proportion of the temporal variation in each domain was aligned along a common underlying temporal dimension. Domain-specific individual components accounted for an additional 19.3%, 7.1%, 17%, and 7.1% of the variance, whereas residual variance accounted for 41.1%, 17.6%, 8.4%, and 17.5%, respectively. The joint component showed consistent loadings across food, MVPA, light exposure, and sleep timing variables, indicating a shared temporal organization across behavioral domains (Table S5). Food timing variables showed generally weaker loadings, consistent with the larger proportion of domain-specific variation observed for this domain.

**Figure 3.**
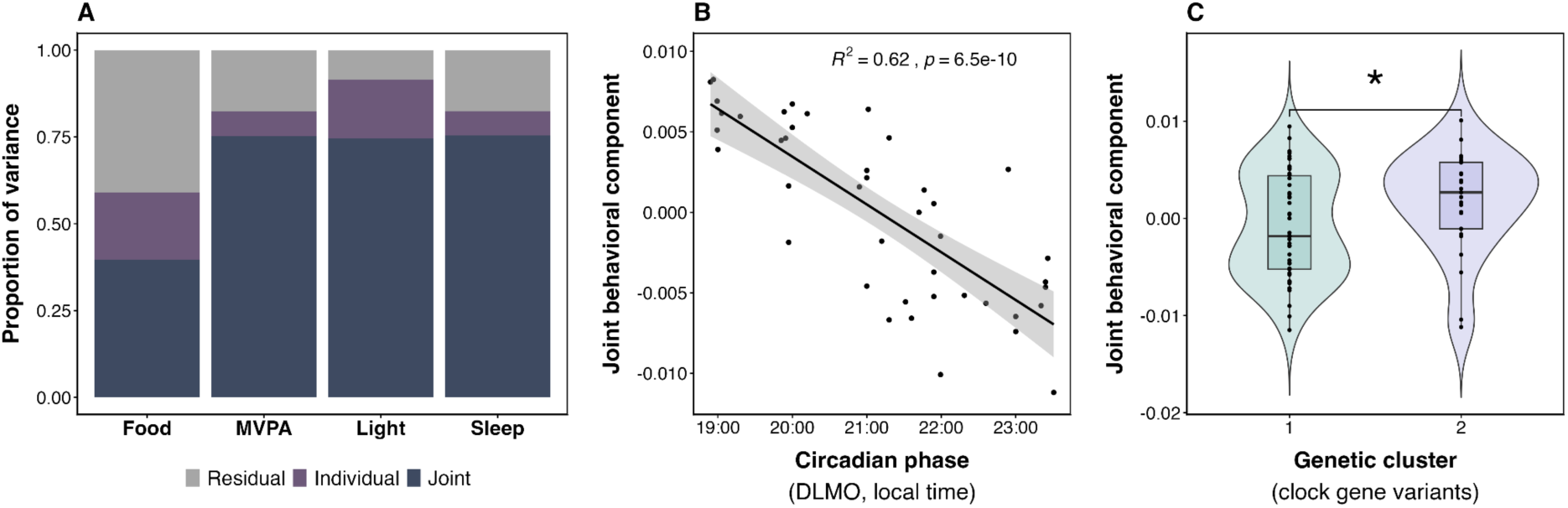
Joint structure across behavioral domains and relationships with circadian and genetic markers. (A) Proportion of variance explained by joint, domain-specific individual, and residual components within each domain (food intake, MVPA, light exposure, and sleep) estimated by JIVE. (B) Correlation between the joint behavioral component and the physiological circadian phase marker DLMO. (C) Comparison between the joint behavioral component and genetic clusters derived from the full set of clock gene variants.

This joint behavioral component captured the temporal distinction between early and late chronotypes, with late chronotypes showing lower values (P < 0.001, r = 0.79). The joint score was also significantly correlated with the physiological circadian phase marker DLMO (Fig. 3B) and with all actigraphy-based circadian phase indicators (Fig. S3). Furthermore, carriers of the T allele of *PER2* rs35333999 variant showed a trend toward lower joint scores compared with non-carriers (n = 8 and n = 67, respectively). Although this difference did not reach statistical significance (P = 0.059, r = 0.22), the direction of the effect was consistent with the overall temporal pattern. Finally, MDS clusters based on clock gene variants exhibited higher values of the joint behavioral component in Cluster 2 (n = 25; enriched for early chronotypes) compared with Cluster 1 (n = 50; enriched for late chronotypes) (P = 0.0499, r = 0.23) (Fig. 3C). Together, these findings reveal a convergence of variation in behavioral timing, circadian phase, and clock genes that defines an integrated temporal phenotype.

## DISCUSSION

Our findings show that, in an underrepresented admixed Latin American population, individuals at the extremes of the chronotype distribution differed in a behavioral component, which closely aligned with circadian phase and was consistent with variation in circadian clock genes, defining an integrated temporal phenotype.

Self-reported early and late chronotypes showed distinct patterns of genetic variation in *BTRC*, with individuals clustering according to chronotype. Interestingly, early chronotypes also showed a slightly higher proportion of observed variants predicted to have deleterious effects. Notably, *BTRC* was among the clock genes harboring the highest number of variants predicted to have deleterious effects in our cohort. *BTRC* encodes an E3 ubiquitin ligase involved in the proteasomal degradation of PER2 (Francisco and Virshup 2024; Ohsaki et al. 2008). Because PER2 stability is a key determinant of circadian period length, alterations in the phosphorylation and ubiquitination pathways regulating PER proteins have long been implicated in chronotype variation (Patke et al. 2020; Takahashi 2017; Von Schantz 2017). Together, these findings highlight *BTRC* as a candidate gene potentially underlying interindividual variation in chronotype in this cohort and warrant further investigation of the functional consequences of BTRC variation on circadian phenotypes. In our study, *PER2* exhibited the highest number of variants per kilobase and the lowest mean PhyloP scores, consistent with reduced evolutionary constraint and a higher tolerance for genetic variation relative to the other circadian genes analyzed. Previous reports have identified signatures of positive selection in *PER2* across geographically diverse populations out of Africa (Cruciani et al. 2008). These observations are consistent with the notion that, while *PER2* harbors substantial genetic diversity, particular variants may have been shaped by population-specific positive selection. Against this background, the *PER2–BTRC* axis may contribute to chronotype variation in this admixed population, providing a rationale for future functional and population-based studies.

The association between the *PER2* rs35333999 variant and activity acrophase under free-living conditions observed in our study is consistent with its previously reported association with intrinsic circadian period measured under forced desynchrony (Chang et al. 2019). Despite the markedly different experimental settings, these independent findings converge to support a role for rs35333999 in human circadian timing. Interestingly, rs35333999 tags an introgressed Neanderthal haplotype displaying a latitudinal allele frequency cline in European populations (Velazquez-Arcelay et al. 2023). Together, the genetic, evolutionary, and circadian evidence identifies rs35333999 as a compelling candidate locus contributing to interindividual variation in human circadian phenotypes.

Physical activity has been recently recognized as a zeitgeber capable of influencing circadian phase (Youngstedt et al. 2019), as was also observed in a cohort of Uruguayan dancers exposed to arbitrary training schedules (Coirolo et al. 2022). At the molecular level, exercise can modulate the expression of core clock genes, with both aerobic and resistance exercise reported to upregulate *BMAL1* and *PER2* expression in skeletal muscle (Shen et al. 2023). On the other hand, the timing of physical activity is a classic behavioral marker of circadian phase (Witting et al. 1990; Halberg et al. 1967). In our study, late chronotypes showed a delay across all activity-timing measures compared with early chronotypes. Moreover, the joint behavioral component accounted for a large proportion of the variance in MVPA timing, indicating strong alignment within a shared temporal organization.

Food timing is aligned to the sleep/wake cycle, and altered patterns are accompanied by rephasing of the transcriptional-translational feedback loop in most peripheral organs (Patton and Hastings 2023, Healy et al 2021). Wehrens et al. (2017) reported that a 5-h delay in meals shifted circadian glucose rhythms and delayed *PER2* expression in white adipose tissue by 1 h, without affecting SCN-driven melatonin and cortisol rhythms. In our analysis, food timing likewise showed greater domain specificity, with a smaller proportion of its temporal variation captured by the joint behavioral component. Late chronotypes had a later timing of last food intake than early chronotypes, exceeding, although consistent with, dinner times previously reported for Uruguayan high school students (Estevan et al. 2020). A south–north European gradient in eating timing has been reported, with the latest food consumption occasion of the day occurring in Spain (Huseinovic et al. 2019). We found no association between chronotype and global ancestry proportions. However, consistent with Uruguay’s demographic history (Sans et al., 1997), Southwestern European ancestry predominated in our sample, highlighting the potential role of cultural factors in shaping population-specific eating timing. In the Río de la Plata region, the consumption of high-caffeine beverages such as yerba mate has been proposed as a contributor to the population’s pronounced eveningness (Tassino & Leone 2025). Notably, late chronotypes in our sample showed later timing of yerba mate and coffee consumption, further supporting the relevance of this cultural context to chronotype variation.

During the Out-of-Africa dispersal, human populations transitioned from relatively stable light/dark cycles to increasingly seasonal and variable photoperiodic environments (Von Schantz 2017; Velazquez-Arcelay et al. 2023). Studies in Brazilian and Eurasian populations (Leocadio-Miguel et al. 2017; Putilov et al. 2019) have suggested a latitudinal cline, with chronotypes becoming progressively later at greater distances from the equator. Interestingly, late chronotypes have been suggested to show greater flexibility in coping with variable light/dark conditions (Zerbini et al. 2021), in line with theoretical predictions from oscillator models (Granada et al. 2013). Uruguay, located between 30° and 35°S, experiences substantial seasonal variation in photoperiod, ranging from approximately 9.8 h in winter to 14.5 h in summer, posing a challenge for circadian entrainment. Time zone alignment may also influence circadian phenotypes (Rodríguez Ferrante and Leone 2024). Although Uruguay lies within the geographic UTC−4 time zone, the country is assigned to UTC−3, resulting in an advance of social time relative to solar time. This discrepancy is further reinforced by cultural practices, such as late evening meals. Together, these multifactorial influences on chronotype make Uruguayan population a valuable model for investigating an integrated temporal phenotype.

A noteworthy finding of this study was the difference in the joint behavioral component between genetic clusters defined by clock gene variation, which also differed in their assignment across chronotypes, reflecting the entangled relationship among endogenous circadian processes and observed temporal organization. The genetic architecture underlying circadian rhythms is highly complex, involving pleiotropic effects, polygenic inheritance, and epigenetic regulation (Burns et al. 2024; Jones et al. 2019a, 2019b; Li et al. 2020; Von Schantz et al. 2021). Although polygenic risk scores for chronotype have been proposed, their applicability across diverse populations remains limited (Biscontin et al. 2025, 2022; Carpena et al. 2025). In contrast, the integration of canonical core circadian genes continues to provide valuable biological insights within multilayered approaches (Nelson et al. 2025). Growing evidence linking chronotype variation to diverse health outcomes has increased interest in circadian typology and has motivated efforts to identify the factors shaping circadian diversity within a translational chronobiology framework (Klerman 2026). Our findings underscore the importance of incorporating multidimensional, population-specific data to better capture the complexity of circadian variation. By integrating behavioral, genetic, and cultural dimensions, our multilayer population characterization provides a framework to advance the understanding of circadian diversity across human populations.

Several limitations should be considered when interpreting these findings. First, the study was based on a relatively small sample for genetic analyses, and therefore the observed associations between clock gene clusters and chronotype should be considered exploratory and require replication in larger cohorts. Second, participants were selected from the extremes of the chronotype distribution, which maximized phenotypic contrast but may limit the generalizability of the findings to individuals with intermediate chronotypes. Third, while sleep, activity, and light exposure were assessed objectively using actigraphy, food intake timing was obtained by self-report and may therefore be subject to reporting bias. Finally, the study was conducted in a sample of young adults from Uruguay, and the extent to which these findings generalize to other age groups or populations remains to be determined.

As recently proposed, chronotypes are embedded within chronotopes (Golombek 2026), where both geography and culture shape circadian rhythms. Our findings support this view by showing that chronotype is expressed as an integrated temporal phenotype spanning behavioral organization, circadian phase, and genetic variation. By extending this framework to an underrepresented admixed southern Latin American population, this study broadens our understanding of the biological and environmental determinants of human circadian diversity.

## METHODS

### Participants

We recruited young adults aged 18–30 years through advertisements posted on Uruguayan university websites, institutional services, and social media platforms between August and October 2024. A total of 361 participants (259 females, 97 males, 4 transgender males, 1 transgender woman; mean age ± SD: 23 ± 3.8 years) completed an online questionnaire assessing sleep, dietary, alcohol consumption, and physical activity habits. Of these, 294 participants (203 females, 87 males; mean age ± SD: 22.9 ± 3.7 years) met the inclusion criteria, which excluded the use of psychotropic, hypnotic, or sleep medications (Fig. 1A). Eligibility for progression to the next study phase required consistency between circadian preference, assessed using the Morningness–Eveningness Questionnaire (MEQ; Horne and Ostberg 1976; Question 19), and chronotype, assessed using the mid-sleep point on free days corrected for sleep debt on weekdays (MSFsc; Roenneberg et al. 2004). Chronotype classification was based on the median MSFsc cutoff (05:04) and restricted to participants within the lower and upper quartiles of the MSFsc distribution (04:09 and 06:04, respectively). A subset of 75 participants (47 females, 28 males, 23.1 ± 3.4 years) continued to the main study (Fig. 1B).

Participants provided DNA oral samples and underwent actigraphy devices during 15-17 days between September and October 2024. The study was evaluated by the Ethics Committee of the School of Psychology, Universidad de la República (Exp. N° 191175-000124-23), and complied with the principles outlined by the Declaration of Helsinki (World Medical Association 2013). Participants signed informed consent.

### Self-report questionnaire

The self-report questionnaire was administered online and included demographic information (age, gender identity, biological sex, birthplace), circadian preference, sleep habits, and timing of first and last coffee, yerba mate infusion (a traditional herbal infusion prepared from *Ilex paraguariensis* leaves containing stimulant compounds), and food consumption on free and work days. Self-reported circadian preference was assessed using Question 19 of the Spanish version of the Morningness–Eveningness Questionnaire (MEQ; Horne and Ostberg 1976): “Do you consider yourself to be?”. Participants classified themselves as “Definitely a morning person”, “More a morning than an evening person”, “More an evening than a morning person”, “Definitely an evening person”, or “Do not know” (Jones et al. 2019a). We assessed sleep habits using the Spanish version of the Munich Chronotype Questionnaire (MCTQ; Roenneberg et al. 2003).

### DNA

We performed genomic DNA sampling using ORAcollect•DNA swabs (DNA-Genotek Inc., Ontario, Canada) for low-pass whole-genome sequencing (0.5X coverage, GRCh37 v6.1, Gencove, Inc., New York, USA). Variant imputation was conducted using a reference panel of 1000Genomas, Simons diversity project and other projects and internal individuals. Concurrence between LPS and WGS Illumina in 30 individuals with high coverage was 99.5% average. Annotation was done with ANNOVAR (Wang et al. 2010). The clock gene set was selected using PLINK (Purcell et al. 2007), and analyses were restricted to variants mapping to these genes. MDS and PCA was performed on both samples after LD-pruning (using parameter-indep-pairwise 50 5 0.2), followed by filtering based on genotype missingness (--geno 0.10), individual missingness (--mind 0.10), and minor allele frequency (--maf 0.01).

The potential functional impact of variants was assessed using multiple in silico prediction algorithms through the Ensemble Variant Predictor (McLaren et al. 2016), including Sorting Intolerant From Tolerant (SIFT; Kumar et al. 2009), Polymorphism Phenotyping (PolyPhen-2; Adzhubei et al. 2010), and Combined Annotation Dependent Depletion (CADD; Kircher et al. 2014), a framework that integrates diverse annotations into a single quantitative score. SIFT classifies variants as “tolerated” or “deleterious”, and may also indicate low-confidence predictions. The algorithm produces a score ranging from 0 to 1, representing the normalized probability that an amino acid substitution is tolerated. Variants with scores < 0.05 are predicted to be deleterious, whereas those with scores ≥ 0.05 are considered tolerated (Ng and Henikoff 2003). PolyPhen-2 assigns variants to the categories “benign”, “possibly damaging”, or “probably damaging”, with scores ranging from 0 to 1. CADD provides a PHRED-scaled score (log-transformed), with higher values indicating greater predicted deleteriousness. The scores reflect the rank of a variant among all possible substitutions in the human genome: 10 corresponds to the top 10% most deleterious variants, 20 to the top 1%, and 30 to the top 0.1% (Rentzsch et al. 2019). PhyloP evolutionary conservation scores were obtained from the UCSC Genome Browser and used to assess nucleotide-level conservation across species (Pollard et al. 2010). Positive scores indicate conservation, whereas negative scores indicate acceleration.

### Clock gene selection

For this exploratory study, we selected 19 canonical circadian clock genes, including *BMAL1, CLOCK, PER1, PER2, PER3, CRY1, CRY2, CSNK1D,* and *CSNK1E* (core transcription–translation feedback loop components); *RORA, RORB, RORC, NR1D1, NR1D2, NFIL3*, and *DBP* (auxiliary regulatory loop components); *FBXL3, FBXL21,* and *BTRC* (mediators of proteasomal degradation of CRY and PER proteins) (Laothamatas et al. 2023).

### Melatonin

We collected hourly saliva samples (1–2 mL, 18:00–01:00) in dim light (<30 lux) on a weekday during the actimetry measure period (Silva et al. 2019; Coirolo et al. 2022). Participants remained in resting position except for brief trips to the bathroom (also dark) and were not allowed to use light-emitting devices during the sampling period. They rinsed their mouths with water before each sample and received a light meal between the 20:00 and 21:00 and between the 22:00 and 23:00 samples. Saliva samples were frozen, stored at 80 C, and later assayed for melatonin using the salivary melatonin competitive ELISA kit from Salimetrics^TM^ (#1-3402) following the manufacturer’s instructions. Individual dim light melatonin onset (DLMO) was estimated using the dlmoR package with the default threshold (Thalji and Spitschan 2026). Melatonin profiles were available for 54 of the 75 participants, of whom 42 yielded valid Hockey-Stick DLMO estimates (21 early and 21 late chronotypes).

### Actigraphy

Participants wore a GeneActive Original (Activinsights, UK) device on their non-dominant wrist for 7–17 consecutive days around the austral spring equinox to minimize seasonal effects. Devices were programmed to record at 10 Hz and provided continuous measures of activity-rest rhythms, physical activity, and light exposure. We processed data with the GGIR package (Migueles et al. 2019) in the RStudio environment. As proxies of circadian phase, we estimated acrophase from parametric cosinor analysis (Cornelissen 2014; Halberg et al. 1967) and the midpoints of the 10-h and 5-h periods with the highest (M10c) and lowest (L5c) accelerometer activity, respectively, using nonparametric methods (Van Someren et al. 1999; Witting et al. 1990).

### Operational definitions

To characterize temporal organization across behavioral and environmental domains, we analyzed the timing of the first and last daily food intake (Wang et al. 2024), the first and last daily moderate-to-vigorous physical activity (MVPA) event (threshold: 93.2 mg; Hildebrand et al. 2014), the first and last daily light exposure above 100 lux (Stone et al. 2020), sleep onset and sleep end. All variables were calculated separately for weekdays and weekends. Food intake timing was obtained by self-report. For actigraphy-derived variables, the first MVPA event was defined as the first bout of MVPA occurring within 5 h after wake-up. The first light exposure event was defined as the first exposure above 100 lux occurring between 1 h before wake-up and 4 h after wake-up. Last MVPA and last light exposure events were defined as the last occurrences within the 4 h preceding sleep onset.

### Data analysis

We conducted all statistical analyses in R (version 4.5.1; R Core Team 2025), within the RStudio environment (version 2025.9.1.401; Posit team 2025). Differences between chronotypes were assessed using the Wilcoxon rank-sum test, and either the chi square test or Fisher’s exact test, as appropriate. Effect sizes were reported as the standardized Wilcoxon effect size (r = Z/√N; R package rstatix; Kassambara 2019) or Cramér’s V (bias-corrected; R package effectsize; Ben-Shachar et al. 2020). P-values were adjusted for multiple comparisons using the Holm–Bonferroni method (Holm 1979). Statistical significance was defined as P ≤ 0.05. A priori power analysis (R package pwr; Champely 2006) indicated that 37 participants per group were required to detect a large effect size (Cohen’s d = 0.76, α= 0.05, power=0.9).

We applied hierarchical clustering on the MDS components obtained from plink to visualize the sample structure for each set of genes. We assessed clustering quality using silhouette analysis (R package cluster; Maechler et al. 2025) which quantifies how similar an observation is to its own cluster relative to other clusters, with values ranging from −1 (poor clustering) to +1 (well-clustered) (Rousseeuw 1987). Nominally significant gene sets (P < 0.05, before Holm adjustment) were further decomposed at the individual gene level to identify potential driver genes.

To assess behavioral temporal organization, we applied Joint and Individual Variation Explained (JIVE; Lock et al. 2013; R package r.jive; O’Connell et al. 2015) to four domains, namely sleep, light exposure, MVPA, and food intake, each represented by four timing variables (first and last event on weekdays and weekends). JIVE decomposes multiblock data into a low-rank joint structure shared across domains, domain-specific individual structures, and residual noise (Di et al. 2019). Ranks were estimated using permutation testing. The individual joint score was used as a composite measure of behavioral temporal organization.

## DATA ACCESS

The data used in this study are not publicly available due to participant privacy and ethical restrictions and are stored locally at Universidad de la República, Montevideo, Uruguay. Aggregated data are available under restricted and controlled access upon reasonable request.

## COMPETING INTEREST STATEMENT

The authors declare no competing interests.

## ACKNOWLEDGMENTS

We specially thank the participants for kindly taking part in this research. We thank Paloma Mombrú and Nicolás Parrillo for their collaboration in data collection, and Federico Garrido, Ana Laura Machado, Rossana Perrone, Ciro Invernizzi, Ivanna Tomasco, Noelle Rivas, for their invaluable help with logistical matters throughout the study. We especially thank the staff of both Facultad de Ciencias and Campus Luisi Janicki, Udelar, for their assistance and access to facilities, and the members of Grupo Cronobiología for their valuable feedback throughout the development of this study. This work was supported by the Comisión Sectorial de Investigación Científica, Programa Grupos I + D [#883158, 2023-2027], Universidad de la República, Uruguay. MM was also funded by Programa de Desarrollo de las Ciencias Básicas (PEDECIBA) and Comisión Académica de Posgrados (CAP, Udelar), Uruguay. Author Contributions: M.M., L.S., A.S., and B.T., conceived and wrote original draft; C.C. and J.C. performed ELISA analysis, L.S., A.S., B.T. supervised the research. All authors have read and agreed to the final version of the manuscript to be published.

## Notes

### Competing Interest Statement

The authors have declared no competing interest.

## REFERENCES

Adzhubei IA, Schmidt S, Peshkin L, Ramensky VE, Gerasimova A, Bork P, Kondrashov AS, Sunyaev SR. 2010. A method and server for predicting damaging missense mutations. Nature methods 7: 248–249.

Albrecht U, ed. 2010. The Circadian Clock. Springer New York, New York, NY.

Arendt J. 2005. Melatonin: Characteristics, Concerns, and Prospects. J Biol Rhythms 20: 291–303.

Ashbrook LH, Krystal AD, Fu Y-H, Ptáček LJ. 2020. Genetics of the human circadian clock and sleep homeostat. Neuropsychopharmacology 45: 45–54.

Ben-Shachar M, Lüdecke D, Makowski D. 2020. effectsize: Estimation of Effect Size Indices and Standardized Parameters. JOSS 5: 2815.

Biscontin A, Russo A, Marnetto D, Pagani L, Costa R, Montagnese S. 2025. Validation and TaqMan Conversion of a Molecular Chronotype Assessment Approach. J Biol Rhythms 40: 19–26.

Biscontin A, Zarantonello L, Russo A, Costa R, Montagnese S. 2022. Toward a Molecular Approach to Chronotype Assessment. J Biol Rhythms 37: 272–282.

Brown SA, Kunz D, Dumas A, Westermark PO, Vanselow K, Tilmann-Wahnschaffe A, Herzel H, Kramer A. 2008. Molecular insights into human daily behavior. Proc Natl Acad Sci USA 105: 1602–1607.

Burns AC, Zellers S, Windred DP, Daghlas I, Sinnott-Armstrong N, Rutter M, Hublin C, Friligkou E, Polimanti R, Phillips AJK, et al. 2024. Sleep inertia drives the association of evening chronotype with psychiatric disorders: epidemiological and genetic evidence. Psychiatry and Clinical Psychology.

Carpena MX, Sanchez-Luquez K, Xavier MO, Santos IS, Matijasevich A, Wendt A, Crochemore-Silva I, Tovo-Rodrigues L. 2025. Accelerometer-derived sleep metrics in adolescents reveal shared genetic influences with obesity and stress in a Brazilian birth cohort study. SLEEP 48: zsae256.

Carvalho FG, Hidalgo MP, Levandovski R. 2014. Differences in circadian patterns between rural and urban populations: An epidemiological study in countryside. Chronobiology International 31: 442–449.

Casiraghi LP, Plano SA, Fernández-Duque E, Valeggia C, Golombek DA, De La Iglesia HO. 2020. Access to electric light is associated with delays of the dim-light melatonin onset in a traditionally hunter-gatherer Toba/Qom community. Journal of Pineal Research 69: e12689.

Champely S. 2006. pwr: Basic Functions for Power Analysis. 1.3–0. https://CRAN.R-project.org/package=pwr.

Chang A-M, Duffy JF, Buxton OM, Lane JM, Aeschbach D, Anderson C, Bjonnes AC, Cain SW, Cohen DA, Frayling TM, et al. 2019. Chronotype Genetic Variant in PER2 is Associated with Intrinsic Circadian Period in Humans. Sci Rep 9: 5350.

Cipolla-Neto J, Amaral FGD. 2018. Melatonin as a Hormone: New Physiological and Clinical Insights. Endocrine Reviews 39: 990–1028.

Coirolo N, Casaravilla C, Tassino B, Silva A. 2022. Evaluation of environmental, social, and behavioral modulations of the circadian phase of dancers trained in shifts. iScience 25: 104676.

Cornelissen G. 2014. Cosinor-based rhythmometry. Theor Biol Med Model 11: 16.

Cox KH, Takahashi JS. 2019. Circadian clock genes and the transcriptional architecture of the clock mechanism. Journal of molecular endocrinology 63: R93–R102.

Cruciani F, Trombetta B, Labuda D, Modiano D, Torroni A, Costa R, Scozzari R. 2008. Genetic diversity patterns at the human clock gene period 2 are suggestive of population-specific positive selection. European Journal of Human Genetics 16: 1526–1534.

Di J, Spira A, Bai J, Urbanek J, Leroux A, Wu M, Resnick S, Simonsick E, Ferrucci L, Schrack J, et al. 2019. Joint and Individual Representation of Domains of Physical Activity, Sleep, and Circadian Rhythmicity. Stat Biosci 11: 371–402.

Dose B, Yalçin M, Dries SP, Relógio A. 2023. TimeTeller for timing health: the potential of circadian medicine to improve performance, prevent disease and optimize treatment. Frontiers in digital health 5: 1157654.

Duffy JF, Rimmer DW, Czeisler CA. 2001. Association of intrinsic circadian period with morningness–eveningness, usual wake time, and circadian phase. Behavioral Neuroscience 115: 895–899.

Egan KJ, Campos Santos H, Beijamini F, Duarte NE, Horimoto ARVR, Taporoski TP, Vallada H, Negrão AB, Krieger JE, Pedrazzoli M, et al. 2017. Amerindian (but not African or European) ancestry is significantly associated with diurnal preference within an admixed Brazilian population. Chronobiology International 34: 269–272.

Emmanuel P, von Schantz M. 2018. Absence of morningness alleles in non-European populations. Chronobiology International 35: 1758–1761.

Estevan I, Silva A, Vetter C, Tassino B. 2020. Short Sleep Duration and Extremely Delayed Chronotypes in Uruguayan Youth: The Role of School Start Times and Social Constraints. J Biol Rhythms 35: 391–404.

Francisco JC, Virshup DM. 2024. Hierarchical and scaffolded phosphorylation of two degrons controls PER2 stability. Journal of Biological Chemistry 300: 107391.

Golombek DA. 2026. From Chronotype to Chronotope. J Biol Rhythms 41: 7–8.

Granada AE, Bordyugov G, Kramer A, Herzel H. 2013. Human chronotypes from a theoretical perspective. PLoS One8: e59464.

Halberg F, Tong YL, Johnson EA. 1967. Circadian System Phase — An Aspect of Temporal Morphology; Procedures and Illustrative Examples. In The Cellular Aspects of Biorhythms (ed. H. Von Mayersbach), pp. 20–48, Springer Berlin Heidelberg, Berlin, Heidelberg.

Healy KL, Morris AR, Liu AC. 2021. Circadian Synchrony: Sleep, Nutrition, and Physical Activity. Front Netw Physiol 1: 732243.

Hildebrand M, Van Hees VT, Hansen BH, Ekelund U. 2014. Age Group Comparability of Raw Accelerometer Output from Wrist- and Hip-Worn Monitors. Medicine & Science in Sports & Exercise 46: 1816–1824.

Holm S. 1979. A simple sequentially rejective multiple test procedure. Scandinavian journal of statistics 65–70.

Horne JA, Ostberg O. 1976. A self-assessment questionnaire to determine morningness-eveningness in human circadian rhythms. Int J Chronobiol 4: 97–110.

Huseinovic E, Winkvist A, Freisling H, Slimani N, Boeing H, Buckland G, Schwingshackl L, Olsen A, Tjønneland A, Stepien M, et al. 2019. Timing of eating across ten European countries – results from the European Prospective Investigation into Cancer and Nutrition (EPIC) calibration study. Public Health Nutr 22: 324–335.

Jones SE, Lane JM, Wood AR, Van Hees VT, Tyrrell J, Beaumont RN, Jeffries AR, Dashti HS, Hillsdon M, Ruth KS, et al. 2019a. Genome-wide association analyses of chronotype in 697,828 individuals provides insights into circadian rhythms. Nat Commun 10: 343.

Jones SE, van Hees VT, Mazzotti DR, Marques-Vidal P, Sabia S, van der Spek A, Dashti HS, Engmann J, Kocevska D, Tyrrell J. 2019b. Genetic studies of accelerometer-based sleep measures yield new insights into human sleep behaviour. Nature communications 10: 1585.

Kassambara A. 2019. rstatix: Pipe-Friendly Framework for Basic Statistical Tests. 0.7.3. https://CRAN.R-project.org/package=rstatix.

Kircher M, Witten DM, Jain P, O’roak BJ, Cooper GM, Shendure J. 2014. A general framework for estimating the relative pathogenicity of human genetic variants. Nature genetics 46: 310–315.

Konopka RJ, Benzer S. 1971. Clock mutants of Drosophila melanogaster. Proceedings of the National Academy of Sciences 68: 2112–2116.

Kumar P, Henikoff S, Ng PC. 2009. Predicting the effects of coding non-synonymous variants on protein function using the SIFT algorithm. Nature protocols 4: 1073–1081.

Lane JM, Qian J, Mignot E, Redline S, Scheer FAJL, Saxena R. 2023. Genetics of circadian rhythms and sleep in human health and disease. Nat Rev Genet 24: 4–20.

Lane JM, Vlasac I, Anderson SG, Kyle SD, Dixon WG, Bechtold DA, Gill S, Little MA, Luik A, Loudon A, et al. 2016. Genome-wide association analysis identifies novel loci for chronotype in 100,420 individuals from the UK Biobank. Nat Commun 7: 10889.

Laothamatas I, Rasmussen ES, Green CB, Takahashi JS. 2023. Metabolic and chemical architecture of the mammalian circadian clock. Cell Chemical Biology.

Leocadio-Miguel MA, Louzada FM, Duarte LL, Areas RP, Alam M, Freire MV, Fontenele-Araujo J, Menna-Barreto L, Pedrazzoli M. 2017. Latitudinal cline of chronotype. Sci Rep 7: 5437.

Li Y, Shan Y, Kilaru GK, Berto S, Wang G-Z, Cox KH, Yoo S-H, Yang S, Konopka G, Takahashi JS. 2020. Epigenetic inheritance of circadian period in clonal cells. eLife 9: e54186.

Liu X, Zhang Y. 2026. Post-Translational Modifications in Animal Circadian Clocks. Advanced Science 13: e21751.

Liu Y, Sancar A. 2025. Biochemical mechanism of the mammalian circadian clock. FEBS letters.

Lock EF, Hoadley KA, Marron JS, Nobel AB. 2013. Joint and individual variation explained (JIVE) for integrated analysis of multiple data types. The annals of applied statistics 7: 523.

Maechler M, Rousseeuw P, Struyf A, Hubert M, Hornik K. 2025. cluster: Cluster Analysis Basics and Extensions. https://CRAN.R-project.org/package=cluster.

McLaren W, Gil L, Hunt SE, Riat HS, Ritchie GR, Thormann A, Flicek P, Cunningham F. 2016. The ensembl variant effect predictor. Genome biology 17: 122.

Migueles JH, Rowlands AV, Huber F, Sabia S, van Hees VT. 2019. GGIR: A Research Community–Driven Open Source R Package for Generating Physical Activity and Sleep Outcomes From Multi-Day Raw Accelerometer Data. Journal for the Measurement of Physical Behaviour 2: 188–196.

Mistlberger RE, Skene DJ. 2005. Nonphotic Entrainment in Humans? J Biol Rhythms 20: 339–352.

Moreno CRC, Vasconcelos S, Marqueze EC, Lowden A, Middleton B, Fischer FM, Louzada FM, Skene DJ. 2015. Sleep patterns in Amazon rubber tappers with and without electric light at home. Sci Rep 5: 14074.

Nelson N, Malhan D, Hesse J, Aboumanify O, Yalçin M, Lüers G, Relógio A. 2025. Comprehensive integrative analysis of circadian rhythms in human saliva. npj Biol Timing Sleep 2: 17.

Ng PC, Henikoff S. 2003. SIFT: Predicting amino acid changes that affect protein function. Nucleic acids research 31: 3812–3814.

O’Connell MJ, Lock EF, Kaplan A, O’Connell MMJ. 2015. Package ‘r. jive.’

Ohsaki K, Oishi K, Kozono Y, Nakayama K, Nakayama KI, Ishida N. 2008. The Role of β-TrCP1 and β-TrCP2 in Circadian Rhythm Generation by Mediating Degradation of Clock Protein PER2. The Journal of Biochemistry 144: 609–618.

Pandi-Perumal SR, Smits M, Spence W, Srinivasan V, Cardinali DP, Lowe AD, Kayumov L. 2007. Dim light melatonin onset (DLMO): A tool for the analysis of circadian phase in human sleep and chronobiological disorders. Progress in Neuro-Psychopharmacology and Biological Psychiatry 31: 1–11.

Paranjpe DA, Sharma VK. 2005. Evolution of temporal order in living organisms. J Circadian Rhythms 3: 7.

Patke A, Young MW, Axelrod S. 2020. Molecular mechanisms and physiological importance of circadian rhythms. Nature reviews Molecular cell biology 21: 67–84.

Patton AP, Hastings MH. 2023. The Mammalian Circadian Time-Keeping System ed. J. Morton. Journal of Huntington’s Disease 12: 91–104.

Paz V, Dashti HS, Burgess S, Garfield V. 2023. Selection of genetic instruments in Mendelian randomisation studies of sleep traits. Sleep Medicine.

Pollard KS, Hubisz MJ, Rosenbloom KR, Siepel A. 2010. Detection of nonneutral substitution rates on mammalian phylogenies. Genome research 20: 110–121.

Posit team. 2025. RStudio: Integrated Development Environment for R. http://www.posit.co/.

Purcell S, Neale B, Todd-Brown K, Thomas L, Ferreira MA, Bender D, Maller J, Sklar P, De Bakker PI, Daly MJ. 2007. PLINK: a tool set for whole-genome association and population-based linkage analyses. The American journal of human genetics 81: 559–575.

Putilov AA, Dorokhov VB, Puchkova AN, Arsenyev GN, Sveshnikov DS. 2019. Genetic-based signatures of the latitudinal differences in chronotype. Biological Rhythm Research 50: 255–271.

R Core Team. 2025. R: A Language and Environment for Statistical Computing. https://www.R-project.org/.

Rentzsch P, Witten D, Cooper GM, Shendure J, Kircher M. 2019. CADD: predicting the deleteriousness of variants throughout the human genome. Nucleic acids research 47: D886–D894.

Rodríguez Ferrante G, Leone MJ. 2024. Solar clock and school start time effects on adolescents’ chronotype and sleep: A review of a gap in the literature. Journal of Sleep Research 33: e13974.

Roenneberg T, Foster RG, Klerman EB. 2022. The circadian system, sleep, and the health/disease balance: a conceptual review. Journal of Sleep Research 31: e13621.

Roenneberg T, Kuehnle T, Pramstaller PP, Ricken J, Havel M, Guth A, Merrow M. 2004. A marker for the end of adolescence. Current Biology 14: R1038–R1039.

Roenneberg T, Wirz-Justice A, Merrow M. 2003. Life between Clocks: Daily Temporal Patterns of Human Chronotypes. J Biol Rhythms 18: 80–90.

Rousseeuw PJ. 1987. Silhouettes: a graphical aid to the interpretation and validation of cluster analysis. Journal of computational and applied mathematics 20: 53–65.

Sans M, Salzano FM, Chakraborty R. 1997. Historical genetics in Uruguay: estimates of biological origins and their problems. Human biology 161–170.

Silva A, Simón D, Pannunzio B, Casaravilla C, Díaz Á, Tassino B. 2019. Chronotype-Dependent Changes in Sleep Habits Associated with Dim Light Melatonin Onset in the Antarctic Summer. Clocks & Sleep 1: 352–366.

Spangenberg L, Fariello MI, Arce D, Illanes G, Greif G, Shin J-Y, Yoo S-K, Seo J-S, Robello C, Kim C, et al. 2021. Indigenous Ancestry and Admixture in the Uruguayan Population. Front Genet 12: 733195.

Stone JE, McGlashan EM, Facer-Childs ER, Cain SW, Phillips AJK. 2020. Accuracy of the GENEActiv Device for Measuring Light Exposure in Sleep and Circadian Research. Clocks & Sleep 2: 143–152.

Takahashi JS. 2021. The 50th anniversary of the Konopka and Benzer 1971 paper in PNAS: “Clock Mutants of *Drosophila melanogaster*.” Proc Natl Acad Sci USA 118: e2110171118.

Takahashi JS. 2017. Transcriptional architecture of the mammalian circadian clock. Nat Rev Genet 18: 164–179.

Tassino B, Leone MJ. 2025. Night owls of Rio de la Plata region: Real-life scenarios to understand the biological clock. Neuroscience 571: 89–95.

Thalji SM, Spitschan M. 2026. *dlmoR* : An Open-Source R Package for the Dim-Light Melatonin Onset (DLMO) Hockey-Stick Method. J Biol Rhythms 41: 301–323.

Van Someren EJ, Swaab DF, Colenda CC, Cohen W, McCall WV, Rosenquist PB. 1999. Bright light therapy: improved sensitivity to its effects on rest-activity rhythms in Alzheimer patients by application of nonparametric methods. Chronobiology international 16: 505–518.

Velazquez-Arcelay K, Colbran LL, McArthur E, Brand CM, Rinker DC, Siemann JK, McMahon DG, Capra JA. 2023. Archaic Introgression Shaped Human Circadian Traits ed. E. Huerta-Sanchez. Genome Biology and Evolution 15: evad203.

Vera B, Dashti HS, Gómez-Abellán P, Hernández-Martínez AM, Esteban A, Scheer FAJL, Saxena R, Garaulet M. 2018. Modifiable lifestyle behaviors, but not a genetic risk score, associate with metabolic syndrome in evening chronotypes. Sci Rep 8: 945.

Von Schantz M. 2017. Natural Variation in Human Clocks. In Advances in Genetics, Vol. 99 of, pp. 73–96, Elsevier.

Von Schantz M, Leocadio-Miguel MA, McCarthy MJ, Papiol S, Landgraf D. 2021. Genomic perspectives on the circadian clock hypothesis of psychiatric disorders. Advances in genetics 107: 153–191.

Wang K, Li M, Hakonarson H. 2010. ANNOVAR: functional annotation of genetic variants from high-throughput sequencing data. Nucleic acids research 38: e164–e164.

Wang L, Chan V, Allman-Farinelli M, Davies A, Wellard-Cole L, Rangan A. 2024. The association between diet quality and chrononutritional patterns in young adults. Eur J Nutr 63: 1271–1281.

Wehrens SMT, Christou S, Isherwood C, Middleton B, Gibbs MA, Archer SN, Skene DJ, Johnston JD. 2017. Meal Timing Regulates the Human Circadian System. Current Biology 27: 1768–1775.e3.

Witting W, Kwa IH, Eikelenboom P, Mirmiran M, Swaab DF. 1990. Alterations in the circadian rest-activity rhythm in aging and Alzheimer’s disease. Biological Psychiatry 27: 563–572.

World Medical Association. 2013. World Medical Association Declaration of Helsinki: ethical principles for medical research involving human subjects. Jama 310: 2191–2194.

Wright KP, McHill AW, Birks BR, Griffin BR, Rusterholz T, Chinoy ED. 2013. Entrainment of the Human Circadian Clock to the Natural Light-Dark Cycle. Current Biology 23: 1554–1558.

Youngstedt SD, Elliott JA, Kripke DF. 2019. Human circadian phase–response curves for exercise. J Physiol 597: 2253–2268.

Zerbini G, Winnebeck EC, Merrow M. 2021. Weekly, seasonal, and chronotype-dependent variation of dim-light melatonin onset. J Pineal Res 70.

